# Antimicrobial stewardship in animal health: experience-based co-design framework to capture progress and accelerate improvement

**DOI:** 10.1101/2025.06.03.657743

**Authors:** KA Hewson, M Binet, K Richards, B Gleeson, S Adsett, G Tuckett, B Kennedy, R Jenner, J Courtice, T Batterham, SW Page, S Britton

## Abstract

Antimicrobial stewardship (AMS) is essential for mitigating antimicrobial resistance (AMR) in animal health but implementing national initiatives is challenging due to diverse influencing factors and the need for context-specific approaches. An Animal AMS Practice Group, comprised of individuals with lived experience overseeing AMS in various animal health contexts was engaged, and through an experience-based co-design approach created a comprehensive AMS Framework that captures progress and supports tangible improvements in AMS practices in each context. The Framework supported a cross-sectoral pilot assessment that helped users identify progress, areas for improvement, and contextualise antimicrobial usage and AMR data, while also motivating further AMS efforts. Despite common barriers to sharing sensitive data, participants willingly shared AMS results for benchmarking and publication. The process demonstrated that co-design coupled with peer learning and expert support are essential to ensuring AMS tools are utilised in practice. Several participants incorporated the Framework into routine practice, with some using it to drive sector-level AMS action. The resulting Framework offers an adaptable, scalable entry point for AMS efforts and a platform for setting meaningful improvement goals. It supports broader opportunities for national-level AMS assessments and strategy development.

**Impact Statement:** This research offers a practical, scalable approach to improving AMS in animal health. By embedding behaviour change into the project through an experience-based co-design process, rather than expecting it to occur afterward, veterinarians, animal managers, and other frontline professionals defined what good AMS looks like in their context and identified achievable actions to improve daily practices that aligned across diverse animal health settings.

The resulting AMS Framework provides a clear, accessible starting point for those overseeing daily AMS activities to benchmark practices, set meaningful goals, and track progress, while also informing national-level prioritisation and decision-making. This user-led model demonstrates that when AMS tools are co-developed with those who will use them and users are empowered to shape the project’s design and outputs, the resulting tools are more likely to be adopted, shared, and sustained.

## Introduction

The global burden of antimicrobial resistance (AMR), where microbes such as bacteria are no longer killed by medicines (e.g. antibiotics) designed to kill or inhibit their growth, is predicted to be the leading cause of human death and a significant burden on healthcare systems (Murray et al. 2022). It is important to recognise that AMR also presents threats to animal health and food security, however the burden of AMR from an animal health perspective globally has been much less the focus of efforts to mitigate AMR, and as such remains largely undefined (Babo Martins et al. 2024; LeJeune 2025). Despite this, there is substantial evidence for AMR resulting in treatment failures and health issues for animals (Bengtsson and Greko 2014; Weese et al. 2015).

‘Antimicrobial stewardship’ (AMS) refers to strategies aimed at preserving the effectiveness of antibiotics and reducing the development of resistance (Weese 2013). Considerable work has been undertaken to develop comprehensive approaches for AMS in practice in human health settings, including assessment frameworks, which are generally designed for health professionals to implement and oversee (Akpan et al. 2016; Australian Commission on Safety and Quality in Health Care 2021; 2023; 2024; Australian Government Aged Care Quality and Safety Commission 2024; Brotherton 2018; Bumpass et al. 2014; Centers for Disease Control and Prevention 2019; Chen et al. 2015; Khanina et al. 2021; Moehring et al. 2017; Monnier et al. 2018; O’Riordan et al. 2021). Due to the complexity of factors influencing antibiotic effectiveness, use and resistance selection, AMS in animal health settings involves continuous improvement in disease prevention, animal husbandry, environmental, infection, prevention and control practices, infection detection, diagnosis and antimicrobial use. While there is much information available on current best practices in these areas, the challenge lies in implementing this AMS evidence into AMS practice by a wide variation of professionals across animal health contexts (Charani and Holmes 2019; Farrell et al. 2023; Golding et al. 2019; King et al. 2018; Norris et al. 2019; Scarborough et al. 2023; Tonne et al. 2023).

Despite substantial description of AMS practices in animal health, the information is often fragmented between general animal health and AMS-specific interventions, narrowly focused (e.g. companion animals only (Antimicrobial Resistance and Stewardship Initiative 2020)), and/or primarily centred on the veterinary prescribing, supply or antimicrobial usage aspects of AMS only. Importantly, those that oversee and influence AMS and antimicrobial usage in animal settings include not only veterinarians but farmers, managers, owners, carers or higher-level staff who require differing approaches to supporting improvements in AMS (Lloyd and Page 2018). Additionally, in commercial farming systems there may be influence from supply chain actors, such as customers, that may have specific requirements that may influence AMS (Singer et al. 2019). This underscores the need for nuanced approaches to improving AMS in animal health settings, as effective change requires a deep understanding of the behaviours in the current system and those with influence within it, and how it can be feasibly adapted without unintended negative consequences (McKernan et al. 2021).

Terms such as ‘patient-centred,’ ‘human-centred,’ ‘co-design,’ ‘participatory approach,’ and ‘community-led approach’ are commonly used in health research to describe direct engagements between researchers, experts, and stakeholders (‘users’) to develop or implement solutions and create change, with approaches varying from gathering insights from users and recommending or enforcing changes, to empowering users to lead and adopt change themselves (Bate and Robert 2006; Donetto et al. 2015; Duea et al. 2022; Robert et al. 2022; Vaughn 2020; Voorberg et al. 2014). The approach selected depends on the extent to which implementation of the solution and practice change are part of the project itself (Australian Healthcare and Hospitals Association 2017; Donetto et al. 2015; Green et al. 2020; Robert et al. 2022; Voorberg et al. 2014).

Projects that aim to work with users to design and implement solutions embed the creation and implementation of change directly within the project, ensuring solutions are actioned and integrated into real-world contexts. For example, experience-based co-design (Bate and Robert 2006), used in healthcare in Australia, creates partnerships between patients and healthcare professionals to co-create and implement solutions that improve patient care. Similarly, Community Engagement (Mitchell et al. 2019; Mitchell et al. 2023; O’Mara-Eves et al. 2015) focuses on collaboratively developing and implementing solutions directly within the unique context for a community. Both approaches move beyond gathering information, identifying issues or providing recommendations for change by fostering active collaboration and taking responsibility for driving change as a defined outcome for the project. Genuine, non-tokenistic engagement of those with lived experience is essential for developing solutions that drive positive, actionable change (Fylan et al. 2021).

Empowering users to actively shape the solution (rather than merely serving as sources of information) ensures the final product is practical, relevant, and tailored to their specific needs and contexts (Voorberg et al. 2014). This, in turn, increases the likelihood that the solution will be used to create long-term, sustainable change at scale, as opposed to being shelved for reference or creating short-term or small-scale benefits (Robert et al. 2022). Further, involving users in the creation process can build their sense of ownership and commitment to the success of the solution, ultimately resulting in them becoming champions for the solution within their sphere of influence well beyond the project’s completion (Fylan et al. 2021). While user-led, implementation-focused co-design approaches have been applied to improve AMS in specific animal health contexts (Currie et al. 2018; Granick et al. 2024; Jelinski et al. 2022; Lloyd and Page 2018; Morgans et al. 2021; van Dijk et al. 2017), a comprehensive approach that serves as a tangible foundation for benchmarking and accelerating action in AMS across all animal health settings remains lacking.

This project aimed to use experience-based co-design to develop a practical and actionable process or tool that is immediately relatable to, and actionable for those overseeing daily AMS practices, to serve as a foundation for reporting and accelerating tangible AMS improvements across any animal health setting. As part of this, the project sought to engage and empower a cohort of individuals who oversee daily AMS practices in diverse animal health contexts with the tools and knowledge needed to drive AMS improvements at the sector level, beyond their own organisations.

## Methods

Ethical approval for this study was obtained from the CSIRO Human Research Ethics committee, reference number 010/24. A co-design process, similar to experience-based co-design in public health settings(Australian Healthcare and Hospitals Association 2017; Green et al. 2020), was utilised for this project and modified as described below.

### Animal AMS Practise Group

Nominations were sought from various sources including livestock industry peak bodies, relevant organisations and animal enterprises, and large corporatised veterinary service providers to identify individuals with direct experience in overseeing AMS. This approach also ensured these stakeholders were aware of the project and could ask questions early in the project. Individuals with lived experience were approached to form an ‘Animal AMS Practise Group’. While representation from every animal health context or sector was desirable, it was unrealistic to expect all sectors to have individuals with the required experience and availability to contribute meaningfully. The focus was on gathering insights and contributions from motivated individuals with practical AMS experience, which was more readily available in some sectors (e.g., livestock and companion animals) than others (e.g., wildlife health management).

Participants were selected based on the following criteria: must have practical oversight of and implement AMS on a daily basis; must have some pre-conceived idea of what they think ‘good’ AMS was; must be motivated to improve their AMS practices; must be linked to relevant AMS networks beyond their daily employment (i.e. influence animal health, biosecurity and/or AMS practices beyond their company through associations, committees etc). Participants could be veterinarians, farmers, facility managers, carers, and animal health professionals.

The Animal AMS Practise Group meetings were designed to foster peer-to-peer learning and included invited expert speakers, with agendas shaped by AMS-related topics of interest to the group members.

### User interviews to identify experience-based AMS elements

Semi-structured interviews were conducted with AMS Practise Group members to gather information regarding their existing AMS programs, along with their thoughts and perspectives on how to identify what ‘improved’ AMS looks like:

- *How is the performance of your antimicrobial stewardship program integrated into daily operations, and assessed? Please specify any benchmarks, indicators or other performance metrics utilised*
- *In ensuring the effectiveness of your AMS program, what methods do you employ for verification and auditing?*
- *Could you provide a detailed description of the process involved in verifying the effectiveness of your AMS program?*

Each interview was recorded and transcribed using Microsoft Teams, and detailed notes were taken with participant permission.

Interviews were also conducted with stakeholders from New South Wales Health and Queensland Health to gather human health perspectives, and additional information from the Responsible Use of Medicines in Agriculture (RUMA 2024) and the Welsh Arwain Defnydd Gwrthficrobaidd Cyfrifol (Arwain DGC 2025) program was captured via email.

The AMS program elements identified from these interviews were used as the basis for creating a comprehensive scope for AMS in animal health settings.

### Literature search to identify AMS elements

A literature search was conducted of journal articles, reports, strategies, and accreditation programs related to building, assessing, and verifying AMS practices in both human and animal sectors. The objective was to identify elements deemed essential for AMS programs or considered ‘good’ AMS practices. Searches were conducted in PubMed and Google Scholar, supplemented by Google searches to identify relevant grey literature, including reports, policy documents, and conference proceedings not indexed in scientific databases. Search terms combined “AMR” and “AMS” (both short and extended forms) with Boolean operators and either “assessment,” “verification,” “indicators,” “metrics,” “benchmarking,” and “auditing”.

Literature was included if it identified ‘AMS’ as a central topic (regardless of definition) and excluded if it focused solely on elements of human medicine without relevance to animal health (e.g. conditions, settings, or procedures not replicated in veterinary medicine) or where the AMS approach or outcomes drew on other existing literature as the source for AMS elements.

The elements comprising AMS identified in published literature were integrated with those gathered from user interviews to create a comprehensive draft scope of AMS elements considered potentially relevant to animal health settings.

### Co-design of an Animal AMS Framework

The list of AMS elements was condensed to remove duplication or grouped where considered highly related (e.g. availability of prescribing guidelines; utilisation of prescribing guidelines). The Animal AMS Practise Group reviewed and discussed the list of AMS areas to determine relevance to their contexts. These discussions resulted in organisation of the elements into a ‘framework’ – a structured approach outlining key elements to guide and support the implementation of AMS. The Animal AMS Practise Group emphasised the need for a flexible yet comprehensive structure that could integrate existing knowledge, capture existing resources, align practices with evidence, identify gaps and advances in AMS progress, and could be adapted to the unique requirements of specific sectors or settings.

The development of the Animal AMS Framework involved iterative stages of review, discussion, feedback, and refinement with the Animal AMS Practise Group as a collective, to ensure the structure and language used was applicable and relevant across all animal health contexts. A second round of interviews was conducted with Animal AMS Practise Group members individually to further refine the framework, identify formats for presentation, and to identify ways in which adaptation to specific contexts would need to occur to facilitate adoption.

### Pilot of the Animal AMS Framework as an assessment tool

The Animal AMS Practise Group sought to use the Framework to informally assess the progress of their individual AMS programs, hence a pilot of the Framework used in this way was conducted.

The Animal AMS Framework elements were organised into a Google form, designed to enable participants to select the description that best reflected the maturity of their current AMS practice, or to choose “other” if none of the options were deemed relevant. When participants selected “other” further exploration was conducted to determine whether this reflected misunderstanding, misinterpretation, or actual irrelevance to their context. This feedback was critical for refining the Framework, ensuring applicability across all contexts and improving language clarity. A free-text section allowed respondents to provide justification of their response and/or feedback on the description, and any other reflections.

The Google form, along with background information on the project, was distributed via email to members of the AMS Practise Group plus five additional individuals with oversight of AMS in animal health contexts who were identified during the initial participant nomination process but were unable to participate in the project due to time constraints or other commitments. The email invited participants to nominate themselves or another representative from their organisation (if relevant) to complete the assessment. Support was offered and provided if requested. Participants were provided three weeks to complete the form between 24 May and 14^th^ June 2024.

All data collected was kept anonymous, although participants could provide their submission date and time if they wished to receive an individualised report that provided their responses benchmarked alongside the anonymised individual responses of other participants.

Feedback was used to finalise a draft of the Framework. The value of the individualised reports was discussed during subsequent Animal AMS Practice Group meetings, along with participant motivations for utilisation of the Framework.

### Comparison of the Animal AMS Framework with Global AMS Studies

The Animal AMS Framework was compared with published studies and policies that identified key AMS areas in animal health to assess its applicability across international contexts. AMS areas outlined by Granick (Granick et al. 2024), Lloyd (Lloyd and Page 2018), Currie (Currie et al. 2018) and the American Veterinary Medical Association (American Veterinary Medical Association 2018) were extracted into an Excel spreadsheet, aligned with elements of the Animal AMS Framework, and analysed for differences and similarities.

## Results

### Animal AMS Practise Group - peer learning and insights

The eleven individuals nominated for the Animal AMS Practise Group who met the requisite criteria were veterinarians or animal health managers from the pig, poultry, feedlot beef, salmon, dairy, and companion animal sectors. Although other individuals, including from different industries, expressed interest, they were unable to commit the required time within the project’s timeframe.

The Animal AMS Practise Group met five times online between February and August 2024 for meetings of approximately 1-2 hours. The breadth of topics the participants wanted to discuss included antimicrobial usage capture and analysis and international activities with potential impact to Australia. Each member also presented an overview of AMS in their context, which fostered peer-to-peer learning. It was found that loose agendas directed by members worked best to allow free-flowing discussion and encourage contributions to the meetings. This approach resulted in extensive discussion and high levels of engagement in the project itself.

A total of 10 interviews were conducted across the pork, feedlot beef, companion animal, dairy, poultry, and salmon sectors to gather insights into participants’ interpretations and applications of AMS.

### Literature review identified hundreds of individual potential AMS elements

The literature review examined 65 articles and reports that met the inclusion criteria for consideration in detail. These were shortlisted to 39 for extraction of AMS elements (Figure 1). 736 AMS elements in total were extracted (547 from human AMS literature; 189 animal AMS literature; Supplementary material, S1) for integration with those gathered from Animal AMR Practise Group insights.

**Figure 1.**
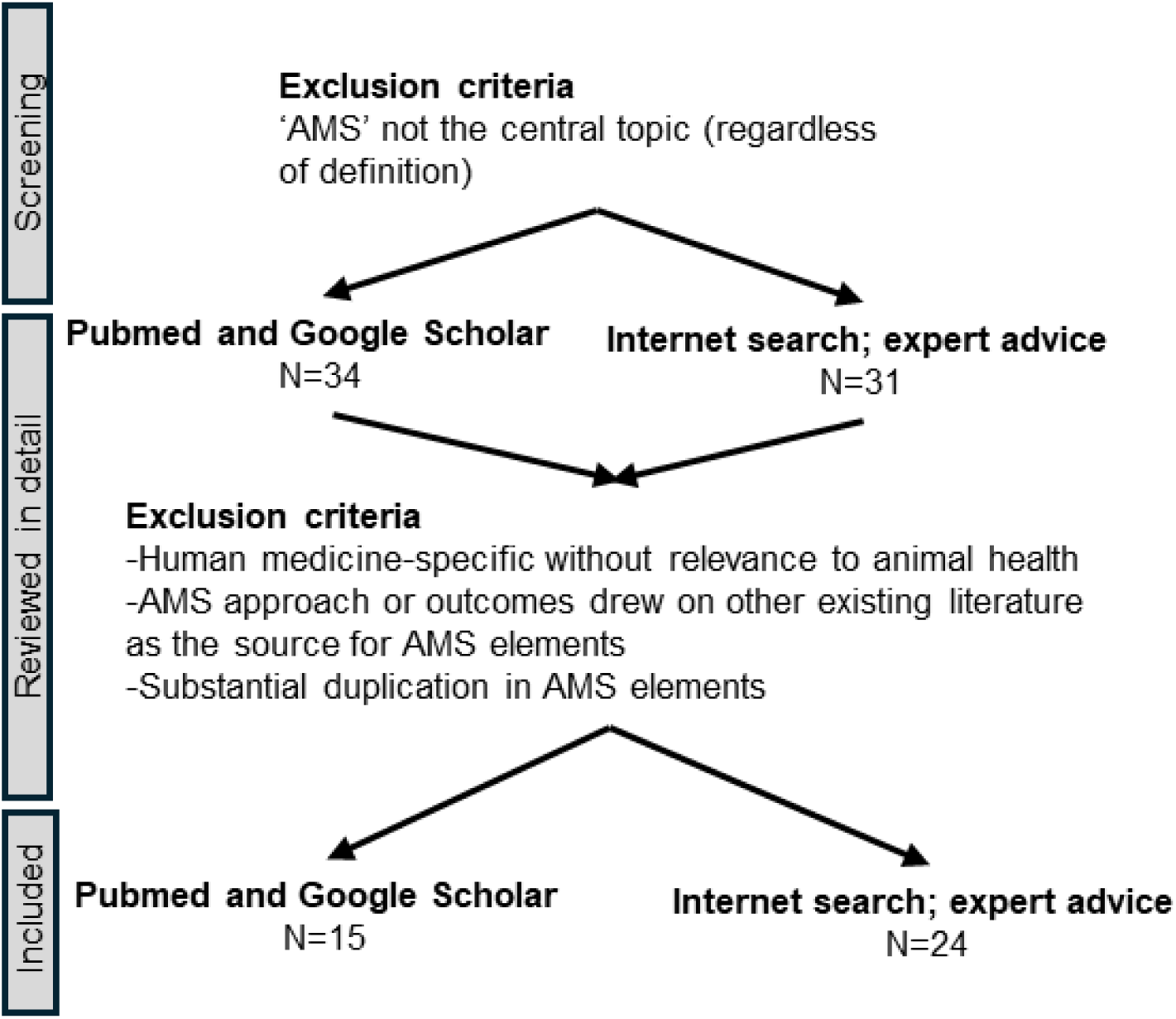
Flowchart detailing screening and selection of articles for extraction of AMS elements that may be relevant in animal health contexts.

### Co-design of an Animal AMS Framework

The process that was followed for co-designing the Animal AMS Framework is outlined in Figure 2. Each co-design process reflects the needs of the individuals involved in the co-design, meaning the process followed in this project would be expected to be generally similar, but not the same, should the process be repeated with a different composition of individuals in the Animal AMS Practise Group.

**Figure 2.**
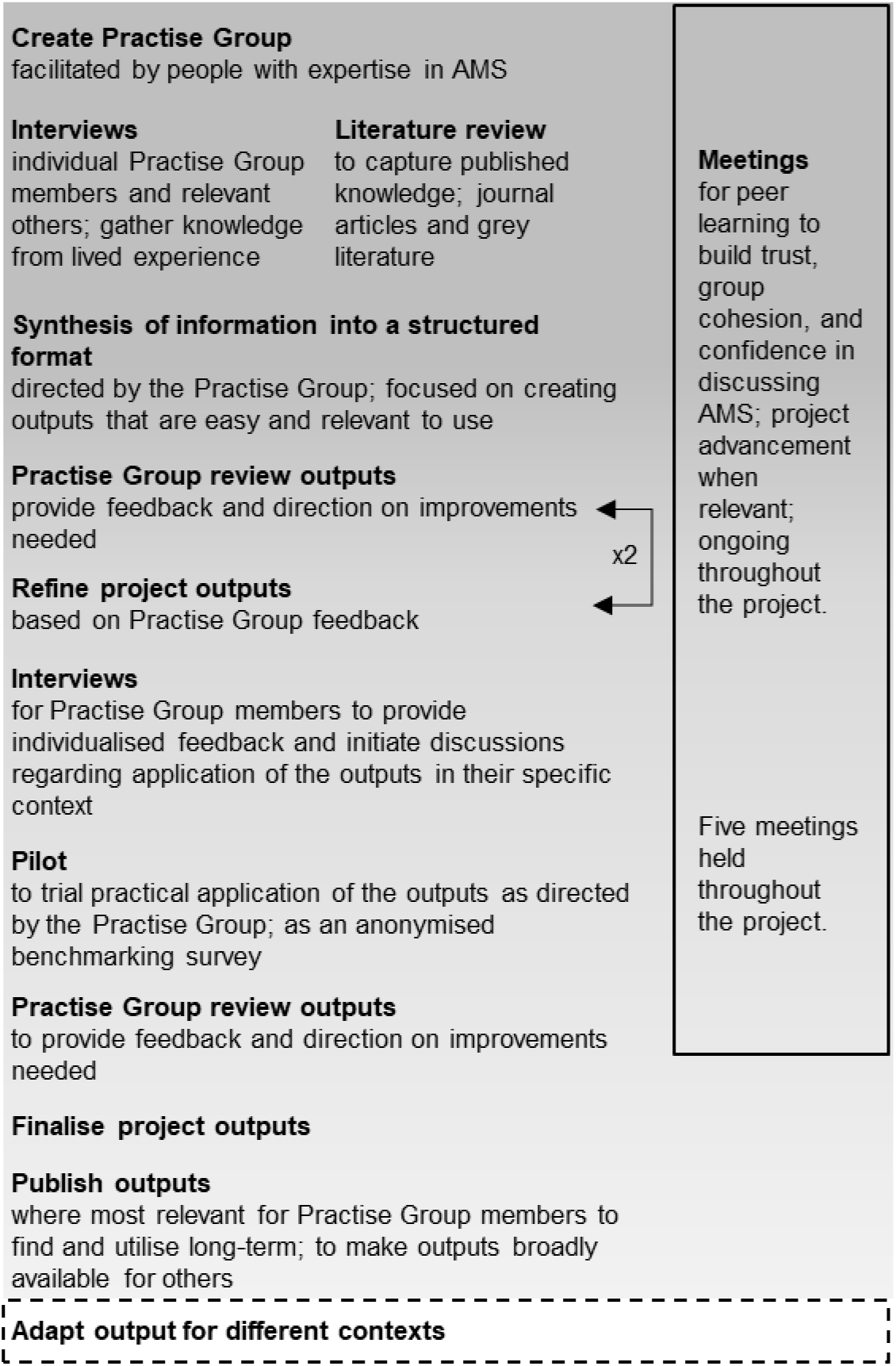
Process implemented during this project as directed by those with lived experience in animal AMS. Dashed box indicates next steps.

After excluding human health-specific AMS elements and combining duplicates or similar elements identified in the literature and the interviews, there were 47 AMS elements grouped into 18 areas that together were considered to describe AMS practices as a whole. No further grouping was agreed upon, and no elements were excluded, disputed, or considered contentious by the Animal AMS Practise Group.

It became clear that many AMS elements were related but represented either a basic or an advanced variation of a practice. As a result, the 18 areas of the Framework each include up to four AMS elements. Initially, the elements under each area were labelled as ‘levels,’ but this terminology implied that the Framework was a compliance tool aiming for the highest ‘level,’ which was not the intended purpose. To better align with the Framework’s goals as a tool to capture and support improvements within context, the language was revised, and the elements categorised as those considered achievable in all contexts (’essential’), or infeasible in many contexts or requiring significant investment to implement (’desirable’). Each ‘desirable’ is considered cumulative in addition to the ‘essential’. The Framework is presented in Table 1.

**Table 1.**
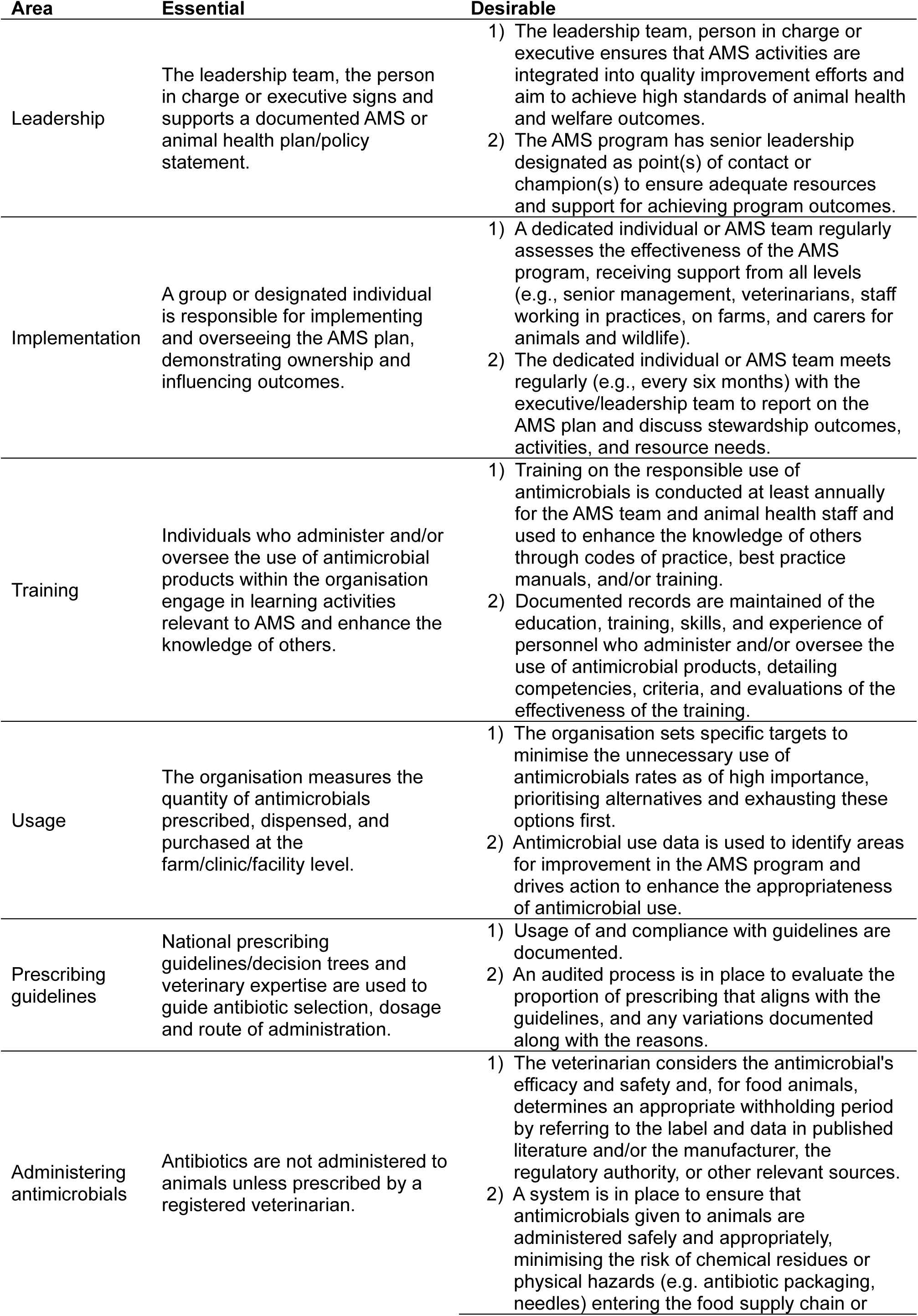
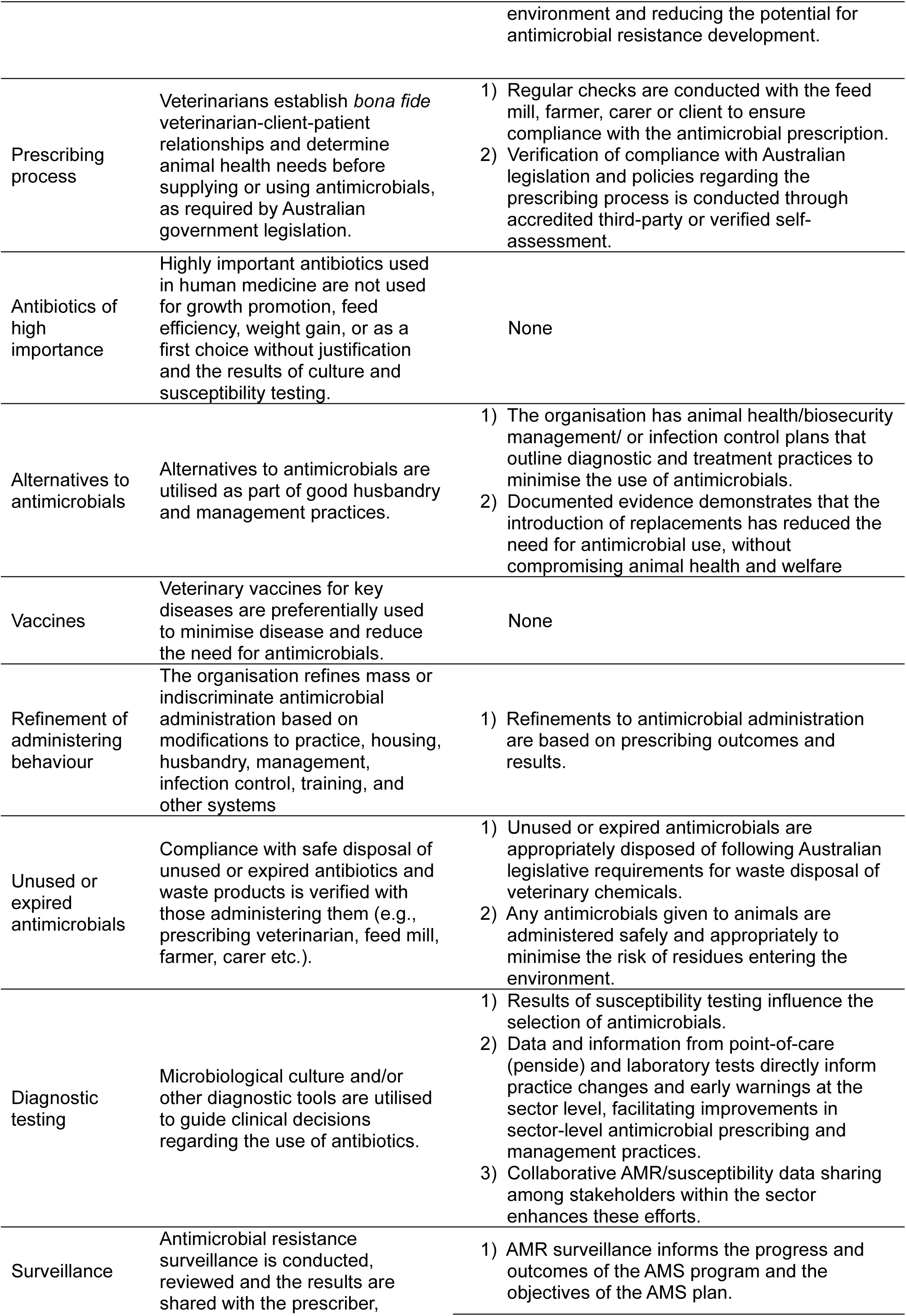
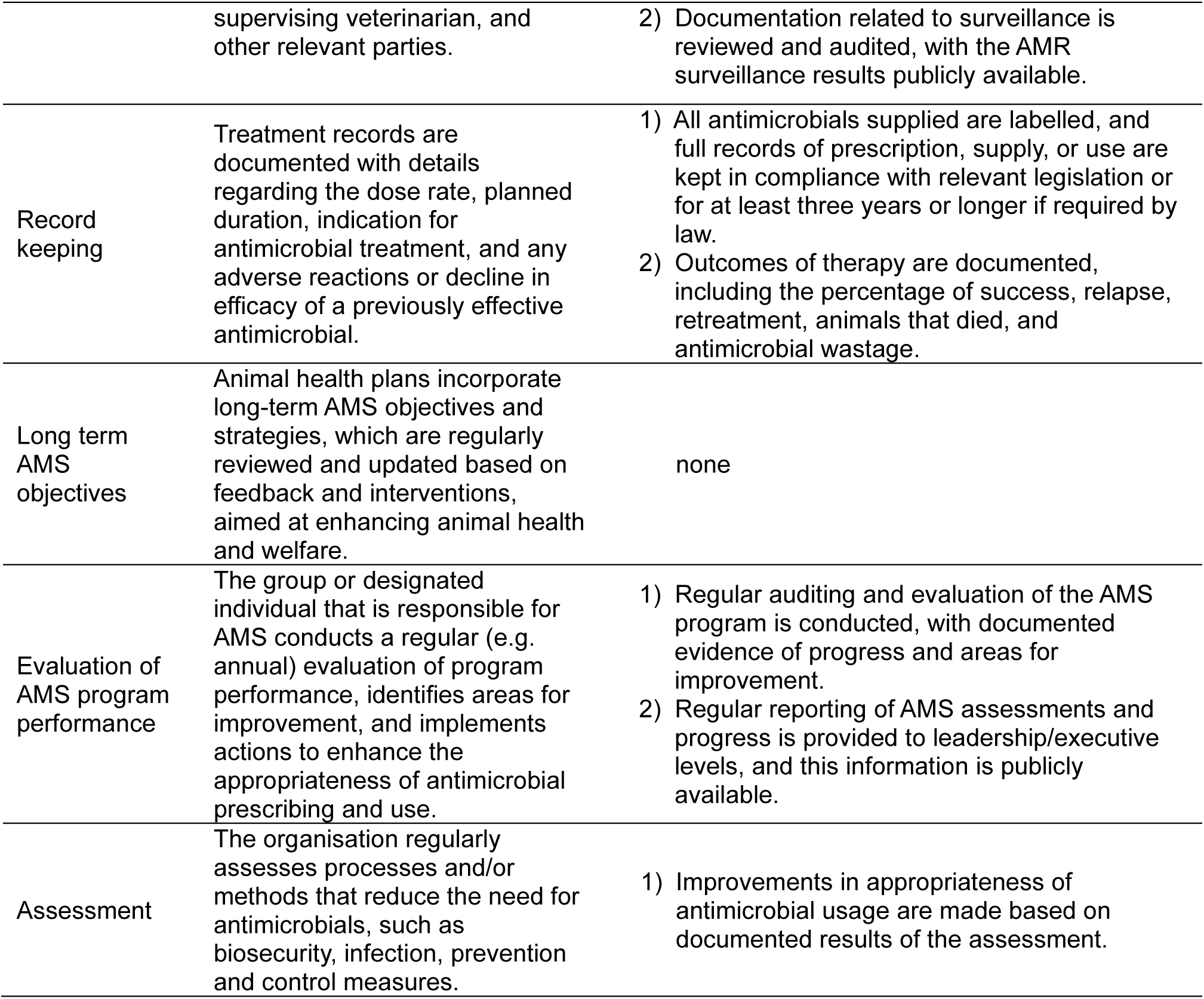
Animal AMS Framework.

Some areas required specifying ‘antibiotics’ (i.e. medicines that inhibit the growth of bacteria specifically) over ‘antimicrobials’ (i.e. collective term for all medicines that inhibit the growth of microbes), however, the majority of the areas focus on antimicrobials broadly. There was considerable discussion as to who was responsible for each element, which influenced the wording as either ‘the responsible person’, ‘group overseeing AMS program’, ‘leadership team’ or ‘organisation’. The terms used throughout the Framework reflect the language and interpretations of those involved in its development. In Australia, the terminology used to describe the importance of antibiotics is ‘high’, ‘medium’ and ‘low’ (Australian Strategic and Technical Advisory Group on Antimicrobial Resistance 2018) and is reflected in the Animal AMS Framework.

### Pilot of the Animal AMS Framework as an assessment tool

#### Use and improvements

Overall, the Animal AMS Framework proved effective as an assessment tool. However, additional discussion and explanation were required to clarify its purpose, benefits, and importance, particularly for those outside the AMS Practise Group who generally needed more guidance and initially viewed the Framework as compliance-focused.

Some Framework elements were less applicable in certain contexts, leading to confusion among participants who were unaccustomed to considering certain aspects of AMS, even when relevant to their context (e.g., companion animal veterinarians needing to consider antimicrobial residues in meat or eggs when treating food-producing animals that are also pets). Ultimately, all elements were deemed relevant after revisions based on pilot feedback.

The layout and ease of use of the pilot AMS assessment was well-received. However, the lack of dashboard functionality in Google Forms specific for what was required resulted in manual handling of responses into Excel for data collation and analysis.

All participants expressed interest in receiving an individualised report benchmarking their assessment against other, anonymised participants. Feedback highlighted the value of individualised reports, a strong interest in future intra- and inter-sector comparative assessments and support for publication of the anonymised assessment outcomes.

Most AMS Practise Group members indicated that they had, or planned to, utilise the Framework as the basis for their organisation’s AMS efforts and requested support from the project team to pursue utilisation of the Framework at a sector level. Their expectation was that the Framework be adapted for utilisation in each context, with adaptations including modifying language and setting targets that broadly resonate with those in the sector, or changing the structure and groupings, without losing the intent of each Framework element.

#### Pilot results

The pilot results were intentionally qualitative, designed to provide a comparative assessment or benchmark among participants. It was not deemed necessary to design the pilot for statistical analysis to identify broader trends as the purpose was to compare the maturity of AMS between pilot participants. Quantitative approaches would be more appropriate for broader intra- or inter-sector assessments, though these would need to be carefully conducted to avoid negative unintended consequences.

Examples of the benchmarking results for each of the areas are presented in Figure 3. These results include those from two animal health professionals from the same company who independently completed the AMS assessment. Their assessments mostly aligned but did include some differing responses. This highlights the potential for an additional consensus-building step in larger companies, AMS teams and within sectors, where individual assessments are followed by discussions to resolve discrepancies and standardise the approach, over time.

**Figure 3.**
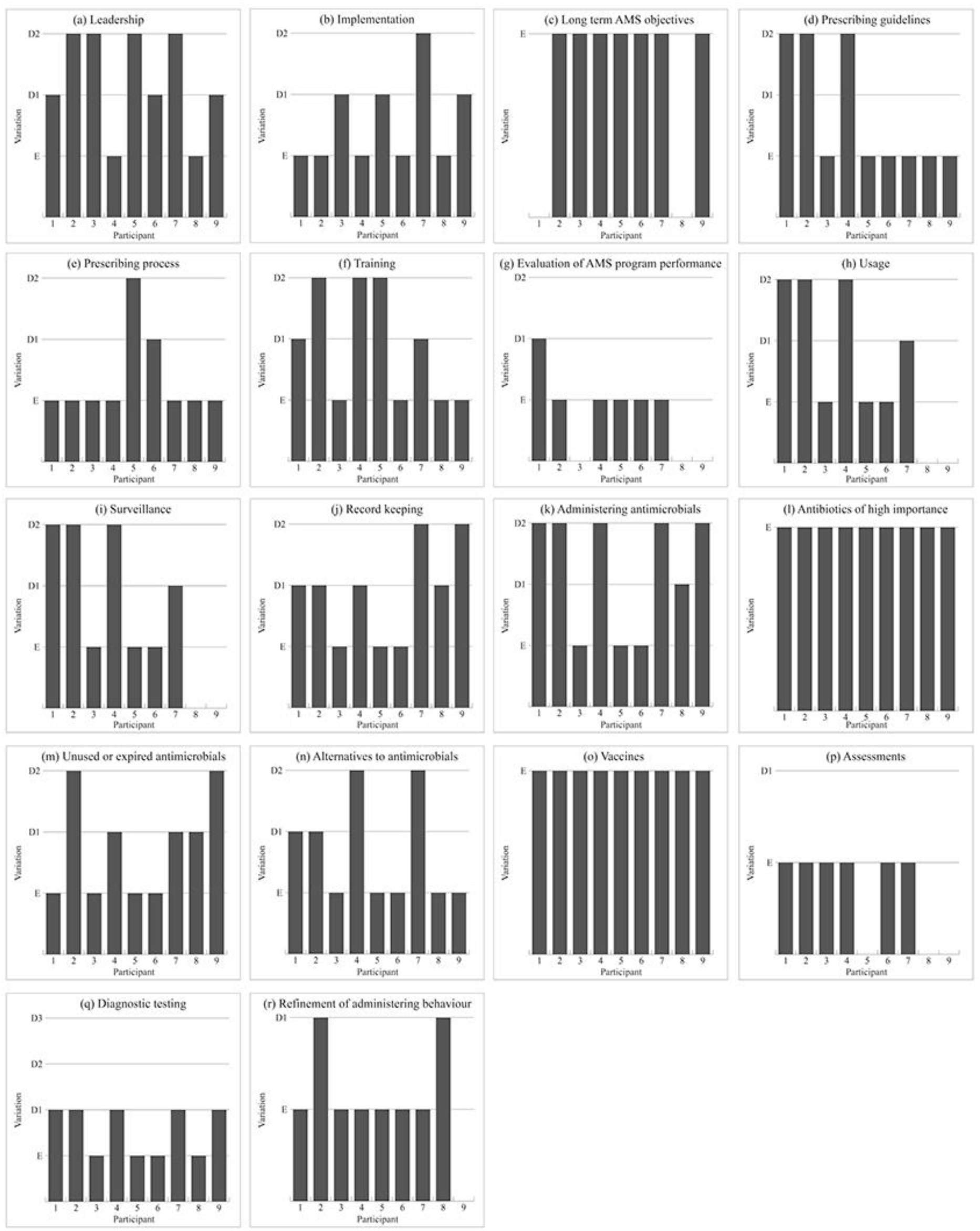
Examples of the benchmarking results for each of the AMS assessment areas. The AMS Framework consists of 47 elements grouped into 18 areas (a – r), with each area containing up to four AMS elements categorised as either “essential,” (y-axis; E) representing achievable and applicable practices across all contexts, or “desirable,” (y-axis; D1 up to D4) which build on the essential elements and often require significant investment or are impractical in many contexts. Each desirable element is cumulative to the essential elements.

The results highlight that for the participants involved in the pilot (i.e. Australian animal health professionals that are motivated to implement and improve AMS programs), there are areas where there is alignment, areas being done well, areas for improvement, and potential inequities or differences in prioritisation of AMS areas by each participant and by different sectors (the latter two likely reflecting their differing contexts). It is important to note that the pilot results are shaped by factors specific to Australia, such as legislation, inequities, supply chain factors and local priorities.

### Comparison of the Animal AMS Framework with Global AMS Studies

The comparison can be found in S2 (Supplementary materials, S2). There were related components between the AMS elements in the Framework and those in the compared publications, however, there was little consistency in alignment across all of them.

The existing publications often combined multiple elements into broad, relatively generic phrases, such as “minimise the selection, maintenance, and dissemination of antimicrobial resistance” or “consistently applying optimal prescribing practices”. The AMS Framework, by comparison, separated out specific, actionable elements, which led to the AMS Framework comprising more elements overall.

All studies compared included these common elements: leadership support and motivation for AMS; accountability for AMS progress through individuals or groups; training and ongoing learning; prescribing guidelines or similar standards; and focus on reducing indiscriminate and mass antimicrobial administration.

Excluding context-specific AMS elements (e.g. client education on AMS; communication with pet owners about AMR) and factors beyond the control of those overseeing daily AMS (e.g., mass media campaigns, laboratory protocols) found in existing publications, the elements identified in the comparison studies that were no reflected in the Animal AMS Framework were ‘consulting technical specialists’ (e.g., microbiologists) and ‘contributing to research’.

The AMS Framework included elements not reflected in the compared publications related to documentation and record-keeping, fostering oversight and accountability through review and verification and following up on whether prescriptions were adhered to. Additionally, the Framework included several elements focused on system-level AMR mitigation and AMS improvements, including publicly sharing AMR data and AMS progress, improving waste management, and sharing disease incidence data.

## Discussion

The Animal AMS Framework encompasses elements of daily AMS and creates opportunities to conduct AMS assessments that capture existing practices. The elements in the Framework tell the AMS story for a particular context, which is informative to understanding factors influencing antimicrobial use and the drivers of AMR, and to identifying meaningful opportunities for improvements, interventions and context-specific targets and benchmarks.

The experience-based co-design approach, centred around the Animal AMS Practise Group comprised of individuals with diverse experiences of AMS and motivation to improve AMS, proved invaluable for several reasons. It helped identify individuals motivated to improve AMS who are also potential influential advocates for sector change; fostered trust between the project team and participants; provided learning opportunities; and enhanced participants’ understanding of AMS and fostered engagement, experiential input and buy-in to ultimately create a behaviour change tool that is relevant, accessible and non-threatening, and therefore more likely to be adopted. All of these factors are critical to successful experience-based co-design (Robert et al. 2022). Additionally, it allowed the project team to recognise and manage biases in their understanding of AMS practices. This was important because developing a tool solely based on literature and expert input would have overlooked the importance of lived experience and context in creating a practical outcome to drive behaviour change. It also would have missed the opportunity to identify advocates for the Framework’s initial adoption and continued advocacy beyond the project’s completion.

Animal AMS Practise Group members became increasingly comfortable and open in discussing AMS during meetings, including traditionally sensitive topics such as antimicrobial usage, setting meaningful targets and the real and perceived risks associated with public release of information. While there is often reluctance to voluntarily release data and information relating to the amount of antimicrobials used, animal health or AMR, participants in the pilot were enthusiastic about sharing their AMS assessment results (Figure 3). This enthusiasm presents significant opportunities to progress efforts to improve AMS and mitigate AMR as the information gathered through the Framework provides context for antimicrobial usage, and may encourage greater willingness to capture, analyse, and report AMU and AMR data and information in ways that are meaningful and useful for progressing AMS at enterprise and sector levels.

When comparing the Framework to existing published animal AMS reviews, it was noted that existing publications often combined multiple elements into broad, general phrases. While these are comprehensive, they lack specific detail, clarity and practicality, making them harder to interpret, apply and verify effectively in daily practice. In contrast, the granular, action-oriented detail of the Framework ensures that it is directly relevant to those implementing AMS on a daily basis. AMS elements from existing publications that were relevant to the scope of the Framework but not integrated by the AMS Practise Group members (i.e. consulting technical experts and contributing to research) may be considered less directly related to daily AMS management, which likely explains their omission. However, these types of elements could be incorporated where relevant during adaptation of the Framework into a specific context, perhaps at a sector-level.

While the pilot and comparison with existing published animal AMS scopes demonstrated that the Framework provides a comprehensive scope of practical AMS activities applicable across all animal health contexts, it does not include the context-specific elements, structure or language necessary for implementation in practice at a sector level. This was evident during the project, as the Animal AMS Practise Group members were unable to agree on further categorisation of the 18 AMS areas, substantial efforts were required to modify the language and terminology to ensure accurate interpretation across diverse animal health contexts, and those in the pilot who were not involved with the development of the Framework were likely to initially misinterpret its purpose. To ensure the Framework is useful at the user level, it is crucial to adapt it to each specific context by incorporating relevant language, targets and element groupings to make it directly applicable and meaningful for those overseeing and implementing AMS in practice. This adaptation step was beyond the scope of this study, however, the suggestion to include an adaptation step indicates that users are seeking a tool that integrates easily into their daily practice. Consequently, with adaptation, the tool is expected to look somewhat different in different contexts, while maintaining the intent of each element.

It was clear that the Framework could not be effectively applied at the user level without support to help users understand how its elements apply to their specific context, including establishing targets and clarifying resourcing limitations. Linking existing tools and experts with those overseeing animal AMS is not new, with existing networks providing connections, expert guidance, and peer learning opportunities (AIAS 2020; Arwain DGC 2025; Bollig et al. 2022; RUMA 2024). This underscores that while the Framework developed in this project serves as a valuable starting point for advancing AMS, adaptation must be coupled with access to trusted, expert support networks for meaningful progress to be made.

It is recommended that adapting the Framework be undertaken in association with literature searches to identify published information relevant to the specific context (e.g., (Bollig et al. 2022) for companion animal AMS) and in consultation with Practise Groups of motivated users. However, a key limitation of this project was restricted access to full-text papers, as many were behind paywalls or required costly subscriptions, which hindered the literature search process for many involved in the project including the Practise Group. This highlights that experts supporting users, or users themselves, are likely to face challenges in accessing all relevant evidence and research needed to inform AMS practice changes which may inhibit adoption of evidence-based solutions.

At a different scale, the Animal AMS Framework offers opportunities for sector- or jurisdiction-level AMS assessments with minimal modification due to its broad applicability. In theory, adapted frameworks should remain aligned with the Framework developed in this project to allow sector- or jurisdiction-level assessments to be conducted using information gathered from the adapted frameworks. This approach would reduce the burden on those overseeing AMS by minimising the need to provide information in multiple formats for various internal and external verification purposes, while also identifying priority areas for investment, training and other strategic support at the sector or jurisdiction level. This would also inform the development or updating of any National AMR Strategy or Action Plan by identifying issues specific to a country. Future work will explore the applicability of the Framework at this scale, and in different countries and AMS settings.

This study had several limitations. The pilot process was cumbersome and error-prone due to the repetitive nature of data handling. A pilot was not originally planned during the project’s design, as the direction depended on how those with lived experience would shape the project through co-design. To streamline future use of the Framework as an assessment tool, an automated system is needed to facilitate comparative analysis and automatically generate individualised reports. This approach could also be expanded for sector- or jurisdiction-level assessments to be undertaken in a way that enables longitudinal analyses.

Future work is needed to evaluate how effectively a focus on improving AMS drives positive practice change and to examine whether AMS assessment results correlate with animal health outcomes and AMR prevalence. Additionally, trained external auditors would be necessary for objective and independent assessments in the long-term. While the current approach of self-evaluation may support progress in internal processes within a specific animal health setting, external assurance or sharing with external stakeholders may require independent assessment to ensure consistency, credibility and reliability.

A follow-up with the AMS Practise Group members over time would be valuable to assess how their participation in the experience-based co-design process has influenced their focus on and improvements in animal AMS, as well as to identify any additional benefits or challenges they experienced. Several Animal AMS Practise Group members have begun adapting the Framework into their daily practices, with some also using it to drive sector-level AMS action independently and beyond the scope of the project, without prompting from the project team. This highlights the value of fully integrating and engaging users into the creation and design of project outputs, empowering them with the knowledge, confidence, and tools needed to become champions for change (Fylan et al. 2021).

## Contributions

K.A.H - Conceptualisation, methodology, investigation, formal analysis, writing - original draft preparation, writing - review and editing, funding acquisition; S.B. - Methodology, investigation, data curation, formal analysis, writing – review and editing; M.B. - Methodology, investigation, project administration; K.R., B.G., S.A., G.T., B.K., R.J., J.C., T.B., M.A. – Methodology, investigation, supervision, validation, writing – review and editing.

## Supporting information

Supplementary data 1 - references

Supplementary data 2 - comparison

## Acknowledgements

We express our appreciation to the livestock industry representatives who took the time to nominate members of their industries to the Animal AMR Practice Group, and those who participated in the pilot and provided valuable feedback.

